# Diversification of Transcription Factor NF-κB in Protists

**DOI:** 10.1101/2021.03.15.435342

**Authors:** Leah M. Williams, Sainetra Sridhar, Jason Samaroo, Ebubechi K. Adindu, Anvitha Addanki, BB522 Molecular Biology Laboratory, Pablo J. Aguirre Carrión, Christopher J. DiRusso, Nahomie Rodriguez-Sastre, Trevor Siggers, Thomas D. Gilmore

## Abstract

In this report, we investigate the evolution of transcription factor NF-κB by examining its structure, activity, and regulation in two protists using phylogenetic, cellular, and biochemical techniques. In *Capsaspora owczarzaki* (*Co*), we find that full-length NF-κB has an N-terminal DNA-binding domain and a C-terminal Ankyrin (ANK) repeat inhibitory domain, and its DNA-binding activity is more similar to metazoan NF-κB rather than Rel proteins. As with mammalian NF-κB proteins, removal of the ANK repeats is required for *Co*-NF-κB to enter the nucleus, bind DNA, and activate transcription. However, C-terminal processing of *Co*-NF-κB is not induced by co-expression of IKK in human cells. Exogenously expressed *Co*-NF-κB localizes to the nucleus in *Co* cells. NF-κB mRNA and DNA-binding levels differ across three life stages of *Capsaspora*, suggesting distinct roles for NF-κB in these life stages. RNA-seq and GO analyses identify possible gene targets and biological functions of *Co*-NF-κB. We also show that three NF-κB-like proteins from the choanoflagellate *Acanthoeca spectabilis* (*As*) all consist of primarily the N-terminal conserved Rel Homology domain sequences of NF-κB, and lack C-terminal ANK repeats. All three *As*-NF-κB proteins constitutively enter the nucleus of human and Co cells, but differ in their DNA-binding and transcriptional activation activities. Furthermore, all three *As*-NF-κB proteins can form heterodimers, indicating that NF-κB diversified into multi-subunit families at least two times during evolution. Overall, these results present the first functional characterization of NF-κB in a taxonomic kingdom other than Animalia and provide information about the evolution and diversification of this biologically important transcription factor.

**Significance:** These results represent the first functional characterization of the biologically important transcription factor NF-κB in a taxonomic kingdom other than Animalia. As such, they provide information on the evolutionary origins and basal diversification of NF-κB outside of metazoans. These results suggest that NF-κB plays life stage-specific roles in *Capsaspora*, the closest unicellular ancestor to all metazoans. Finally, the analysis of three NF-κB proteins in a single choanoflagellate indicates that choanoflagellates have subclasses of NF-κBs, which can form heterodimers, suggesting that NF-κB subunit expansion and diversification has occurred at least twice in evolution.

## Introduction

Transcription factor NF-κB (Nuclear Factor-κB) has been extensively studied for its roles in development and immunity in animals from sponges to humans (1–3). Only within the last few years has it been discovered that NF-κB’s appearance pre-dates metazoan life; that is, that certain single-cell eukaryotes, namely some choanoflagellates and the holozoan *Capsaspora owczarzaki*, also contain genes encoding NF-κB-like proteins (4, 5). In this paper, we present the first functional characterization of this important transcription factor in single-celled protists.

Protists comprise a large group of eukaryotes that are either unicellular or multicellular with poorly differentiated tissue, and they make up one of the six major Kingdoms (6). Presumably, protists have a common ancestor, but they are now known to be an extensively diverse collection of organisms with several major supergroups. Two protists that have been studied reasonably well are *Capsaspora* and the taxonomic class of choanoflagellates, both of which are in the Opisthokonta subgroup.

*Capsaspora* is a single-celled eukaryote that is thought to be among the closest unicellular relatives to animals (i.e., basal to sponges) and is the sister group to the Filozoa (the clade comprising metazoans and choanoflagellates) (7). *Capsaspora* was originally discovered as an amoeba-like symbiont in the hemolymph of the fresh-water snail *Biomphalaria glabrata. Capsaspora* kills sporocysts of the flatworm *Schistosoma mansoli*, the causative agent of schistosomiasis in humans, which also inhabits *B. glabrata* (8). More recently, the life cycle of *Capsaspora* has been shown to contain three different cell configurations (8). Under *in vitro* culture conditions, *Capsaspora* grow primarily as filopodial cells, which attach to the substrate and undergo active replication until the end of the exponential growth phase. Then, cells start to detach, retracting their branching filopodia and encysting. During this cystic phase, cell division is stopped. Alternatively, filopodia cells can form a multicellular aggregative structure by secreting an unstructured extracellular matrix that promotes aggregation but prevents direct cell-cell contact (9). The genome of *Capsaspora* contains many genes involved in metazoan multicellular processes including integrins, protein tyrosine kinases, and transcription factors, including NF-κB (4). Furthermore, RNA-sequencing has revealed that each life stage contains distinct transcriptomic profiles (9). However, the details of how these life-stage processes and transitions are carried out on the molecular level in a unicellular organismal context are still unclear.

A second class of protists of interest for evolutionary biologists includes the choanoflagellates. These flagellated eukaryotes comprise over 125 species of free-living unicellular and colonial organisms distributed in nearly every aquatic environment (10). Choanoflagellates are also widely regarded as being close living relatives to the animals, and they are capable of asexual and sexual reproduction (10). The feeding of choanoflagellates on bacteria provides a critical ecological role within the global carbon cycle by linking trophic levels. Until recently, little was known about the genomic diversity of choanoflagellates, with only two published genomes of *Monosiga brevicollis* and *Salpingoeca rosetta* (11, 12). Neither of these species contains homologs to NF-κB, and it was thought for nearly a decade that this transcription factor had been lost in the evolutionary branch containing choanoflagellates. However, in 2018, Richter et al. (5) reported the transcriptomes of 19 additional choanoflagellates, and 12 of these choanoflagellates expressed NF-κB-like genes, and several of these species that contained multiple NF-κB-like transcripts. Amazingly, sequence comparisons have revealed that choanoflagellates are generally as genetically distant from each other as a mouse is from a sea sponge, a testimony to the modern day diversity among this taxa (5).

Proteins in the NF-κB superfamily are related by a conserved N-terminal Rel Homology Domain (RHD) containing sequences important for dimerization, DNA binding, and nuclear localization (1). All NF-κB proteins bind to a collection of related DNA sites known as κB sites, and NF-κB proteins do so as either homodimers or heterodimers. In animals from insects to humans, there are two subgroups of NF-κB proteins that differ in their C-terminal domain sequences and DNA-binding site preferences (1). One subgroup includes the traditional NF-κB proteins (p52/p100, p50/p105, Relish) that contain C-terminal inhibitory repeats known as Ankyrin repeats (ANK), whereas the second group consists of the Rel proteins (RelA, RelB, cRel, Dorsal, Dif) that contain C-terminal transactivation domains. Among basal organisms---including cnidarians, poriferans, and some protists---only NF-κB-like proteins have been found (3). Indeed, no Rel proteins have been identified in any organism basal to flies.

In most metazoans, the activity of NF-κB proteins is controlled by subcellular localization, wherein an inactive NF-κB dimer is sequestered in the cytoplasm due to interaction with inhibitory IκB sequences (including the C-terminal ANK repeats of NF-κB proteins). Many upstream signals, including the binding of various ligands to conserved upstream receptors (e.g., Toll-like Receptors (TLRs), Interluekin-1 receptors (IL-1Rs), and tumor necrosis factor receptors (TNFRs)), lead to the initiation of a signal transduction pathway culminating in activation and nuclear translocation of NF-κB (1, 2). In the non-canonical pathway, the translocation of NF-κB from the cytoplasm to the nucleus is initiated through the phosphorylation of serine residues C-terminal to the ANK repeats, which leads to removal of the C-terminal ANK repeats by a proteasomal processing that begins at the C terminus and stops within a glycine-rich region (GRR) near the end of the RHD (13). Relieved of inhibition, the NF-κB dimer is able to translocate to the nucleus, bind DNA, and activate its transcriptional targets for a given biological outcome.

Transcriptomic and genomic sequencing has revealed that NF-κB and homologs of many of its upstream regulators are present in most eukaryotes from protists to vertebrates (3, 14). However, the numbers and structures of these signaling proteins vary across species, and generally become more complex and numerous through evolutionary time (14). Furthermore, in the most basal groups of metazoans (cnidarians and sponges), only single homologs to NF-κB exist within their genomes (3).

Herein, we have characterized molecular functions of transcription factor NF-κB in two unicellular protists using phylogenetic, cellular, and biochemical techniques. We find that like the human p100 protein, some unicellular NF-κB proteins require removal of C-terminal ANK repeats to enter the nucleus and bind DNA. However, *Co*-NF-κB does not undergo IKK-mediated processing, and homologs to IKK do not exist in *Capsaspora* or choanoflagellates. Furthermore, we show that the multiple NF-κBs of a single choanoflagellate can form heterodimers, a first finding in an organism outside of the kingdom Animalia, suggesting that choanoflagellates contain their own subclasses of NF-κBs, much like in vertebrates and flies. These results are the first functional characterization of NF-κB in a taxonomic kingdom other than Animalia.

## Results

### Protist NF-κB proteins vary in domain structure and choanoflagellates show evidence of gene duplication

Suga et al. (4) reported the presence of a single gene encoding an NF-κB-like protein in *C. owczarzaki* (*Co*). The overall protein structure of *Co*-NF-κB is similar to most other basal NF-κB proteins known to date, in that it has an N-terminal RHD, followed by a glycine-rich region (GRR), and five C-terminal ANK repeats (15–18). However, *Co*-NF-κB is larger than other NF-κB homologs, due primarily to additional residues C-terminal to the ANK repeats (Fig. 1A), Recently, Richter et al. (5) showed that the transcriptomes of several choanoflagellates had NF-κB-like genes, even though NF-κB-like genes are not present in two commonly studied choanoflagellates (*M. brevicollis* and *S. rosetta*) (11, 12). Overall, 12 of the 21 choanoflagellates are now known to contain NF-κB-like genes, and among those 12, there are one to three NF-κB transcripts present. A phylogenetic comparison suggests that many of these NF-κBs arose from gene duplications within a given species because the multiple NF-κBs from a given species often cluster closely to each other (for example, *Diaphanoeca grandis* and *Salpingoeca helianthica*) (Fig. 1B). Nevertheless, there are some choanoflagellates that have multiple NF-κBs that cluster separately with the NF-κBs of other choanoflagellate (e.g., *Acanthoeca spectabilis* and *Savillea parva*). In contrast to what is seen in most basal metazoans, these choanoflagellates express transcripts that primarily encode RHD sequences, with no C-terminal GRRs or ANK repeats. However, some choanoflagellate NF-κBs do contain extended N-termini with homology to sequences not normally associated with NF-κBs in vertebrates (Fig. 1A, pink bar).

**Fig. 1.**
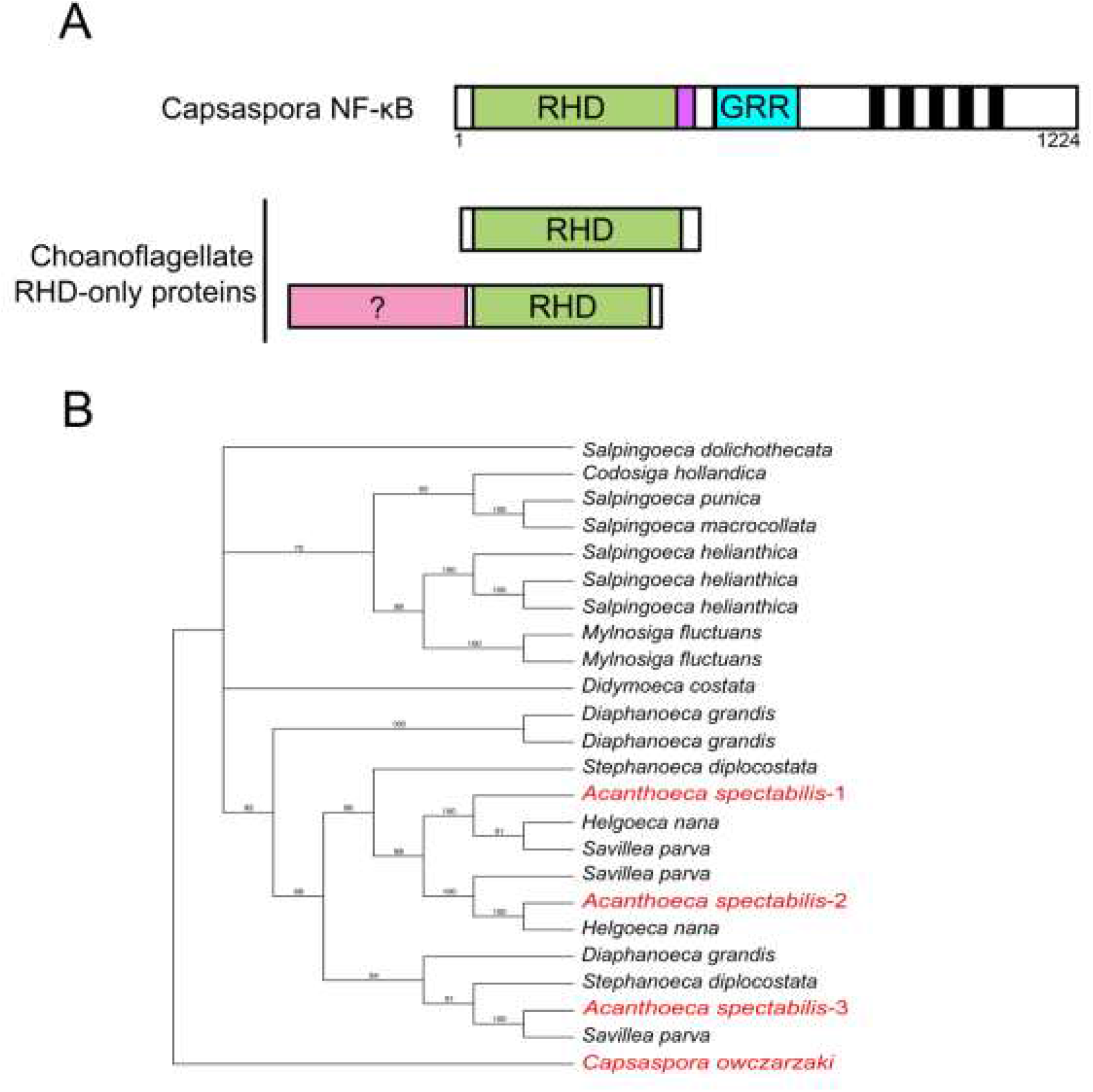
Protist NF-κB proteins differ in domain structure and choanoflagellates show evidence of gene duplication. (A) The general domain structures of both *Capsaspora* NF-κB and the choanoflagellate RHD-only proteins. Green, RHD (Rel Homology Domain); Purple, nuclear localization sequence; Blue, GRR (gylcine-rich region); Black bars, Ankryin repeats; Pink, sequences in choanoflagellates that are not typically seen in other organisms. (B) A phylogenetic estimation using maximum likelihood (bootstrapped 1000 times) of choanoflagellate and *Capsaspora* RHDs. The NF-κB proteins used in these studies are highlighted in red.

### DNA binding, nuclear translocation, and transactivation by Co-NF-κB

To investigate the overall DNA binding-site specificity of *Co*-NF-κB, we first characterized the activity of a bacterially expressed *Co*-NF-κB RHD-only protein by protein binding microarray (PBM) analysis on an array containing 2592 κB-like sites and 1159 random background sequences (for array probe sequences, see (17)). By comparison of the z-scores for binding to DNA sites on the PBM, the DNA-binding profile of *Co*-NF-κB is most similar to NF-κBs from the sea anemone *N. vectensis* and human p52, and it is distinct from human c-Rel and RelA (Fig. 2A).

**Fig. 2.**
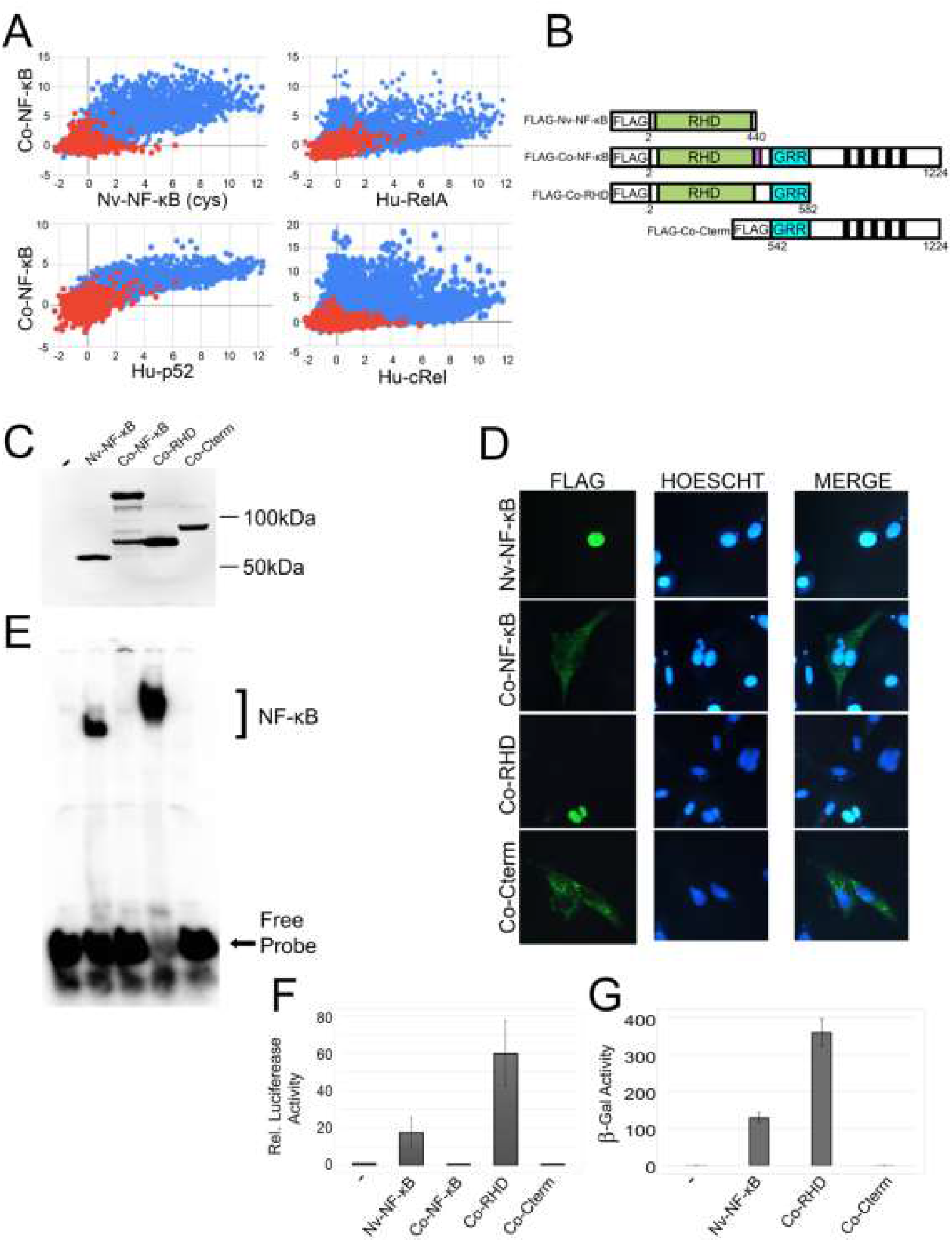
DNA-binding and transcriptional activation activity of *Co*-NF-κB. (A) Protein binding microarray (PBM) DNA-binding profiles of Co-NF-κB as compared to *Nematostella vectensis* (*Nv*) NF-κB cysteine (cys) allele (top, left), human (Hu) RelA (top, right), human p52 (bottom, left), and human cRel (bottom, right). The axes are z-scores. Red dots represent random background sequences, and blue dots represent NF-κB binding sites. (B) FLAG-tagged expression vectors used in these experiments. From top to bottom, the drawings depict the naturally shortened *Nv*-NF-κB, the full-length *Co*-NF-κB protein, an N-terminal-only mutant containing the RHD and GRR (*Co*-RHD), and a C-terminal-only mutant containing the ANK repeats and other C-terminal sequences (*Co*-Cterm). (C) Anti-FLAG Western blot of lysates of HEK 293T cells transfected with the indicated expression vectors. (D) Indirect immunofluorescence of DF-1 chicken fibroblast cells transfected with the indicated expression vectors. Cells were then stained with anti-FLAG antiserum (left panels) and HOESCHT (middle panels), and then MERGED on the right panels. (E) A κB-site electromobility shift assay (EMSA) using each of the indicated lysates from (C). The NF-κB complexes and free probe are indicated by arrows. (F) A κB-site luciferase reporter gene assay was performed with the indicated proteins in HEK 293 cells. Luciferase activity is relative (Rel.) to that seen with the empty vector control (1.0), and values are averages of three assays performed with triplicate samples with standard error. (G) A GAL4-site *LacZ* reporter gene assay was performed in yeast Y190 cells. Values are average units from seven assays performed with duplicate samples with standard error.

To investigate properties of *Co*-NF-κB in cells, we created pcDNA-FLAG vectors for full-length *Co*-NF-κB and two truncation mutants, one (FLAG-*Co*-RHD) containing the N-terminal RHD sequences including the NLS and the GRR, and a second (FLAG-*Co*-Cterm) consisting of the C-terminal ANK repeat sequences and downstream residues (Fig. 2B). As a control, we used the active, naturally truncated *N. vectensis* (*Nv*) FLAG-tagged *Nv*-NF-κB protein that we have characterized previously (19) (Fig. 2B). As shown by anti-FLAG Western blotting, each plasmid expressed a protein of the appropriate size when transfected into HEK 293T cells (Fig. 2C).

In sponge and some cnidarian NF-κBs, removal of C-terminal ANK-repeat sequences are required for nuclear localization in vertebrate cell-based assays (15–17). Based on those results, we next transfected each FLAG expression plasmid into DF-1 chicken fibroblast cells and performed indirect immunofluorescence using anti-FLAG antiserum (Fig. 2D; Supplemental Table 1). Full-length *Co*-NF-κB and *Co*-Cterm were both located primarily in the cytoplasm of these cells (99.9% and 94%, respectively). In contrast, the *Co*-RHD and control *Nv*-NF-κB proteins were both primarily nuclear, as evidenced by co-localization with the Hoechst-stained nuclei (Fig. 2D; Supplemental Table 1). Thus, the removal of the ANK repeats allows *Co*-NF-κB to enter the nucleus, consistent with what is seen with other metazoan RHD-ANK bipartite NF-κB proteins.

To further assess the DNA-binding activity of *Co*-NF-κB proteins, whole-cell extracts from 293T cells transfected with each of the FLAG constructs were analyzed in an electrophoretic mobility shift assay (EMSA) using a high affinity κB-site probe. Extracts containing overexpressed *Nv*-NF-κB and *Co*-RHD bound the κB site avidly, whereas extracts containing full-length *Co*-NF-κB and *Co*-Cterm showed essentially no κB site-binding activity (Fig. 2E).

We also assessed the ability of *Co*-NF-κB proteins to activate transcription in reporter gene assays in HEK 293 cells using a κB-site reporter. *Co*-RHD and *Nv*-NF-κB activated transcription well above control levels (i.e., *Co*-RHD was ~60-fold above the negative control; Fig. 2F). In contrast, full-length or Cterm *Co*-NF-κB proteins showed little to no ability to activate transcription. From these data, the ability to activate transcription of a κB site gene locus appears to be a property of sequences within the N-terminal half of *Co*-NF-κB. We also assessed the ability of *Co*-RHD to activate transcription in reporter gene assays in yeast cells using a GAL4-site reporter. Indeed, the N-terminal half (RHD) of *Co*-NF-κB activated transcription strongly, nearly 1000-fold above the GAL4 (aa 1-147) alone negative control. The transactivation ability of the GAL4-RHD sequences of *Co*-NF-κB in yeast suggests that this is an intrinsic property of these sequences.

Taken together, the results in this section show that *Co*-RHD can bind DNA, activate transcription, and localizes primarily to the nucleus, unlike the inactive full-length *Co*-NF-κB protein, consistent with findings with most NF-κBs from sponges to humans when assayed in vertebrate cells (15–17, 19).

### IKK-mediated processing of NF-κB appears to have evolved with the rise of multicellularity

As we describe above for *Co*-NF-κB, vertebrate NF-κB p100 requires the removal of its C-terminal ANK repeats to enter the nucleus and activate transcription (13). This proteasome-mediated processing of p100 is initiated by phosphorylation of a C-terminal cluster of serine residues by an IκB kinase (IKK) (13). We have previously shown that some basal organisms, including NF-κB proteins from two cnidarians and one sponge, contain homologous C-terminal serines that can be phosphorylated by IKKs to initiate proteasome-mediated processing in human cell culture assays (15–17). Examination of the C-terminal sequences of *Co*-NF-κB failed to identify any C-terminal serine clusters similar to other NF-κBs that undergo IKK-initiated processing. Nevertheless, we performed a series of experiments that examined the ability of IKK to induce processing of *Co*-NF-κB by co-transfecting HEK 293T cells with *Co*-NF-κB and several IKK proteins, including two from humans and one from a sea anemone (15–17). In all cases, co-expression of the IKK did not induce processing of *Co*-NF-κB (Fig. 3A), beyond the small amount of constitutive processing of *Co*-NF-κB that occurs even in the absence of IKK (Fig. 3A and Fig. 2C). Of note, the lower *Co*-NF-κB band seen in these extracts was roughly the same size as the predicted RHD (Fig. 2C), and incubation of transfected cells with the proteasome inhibitor MG132 reduced the appearance of the lower band, suggesting that it arises by proteasomal processing of full-length *Co*-NF-κB (Supplemental Fig. 1).

**Fig. 3.**
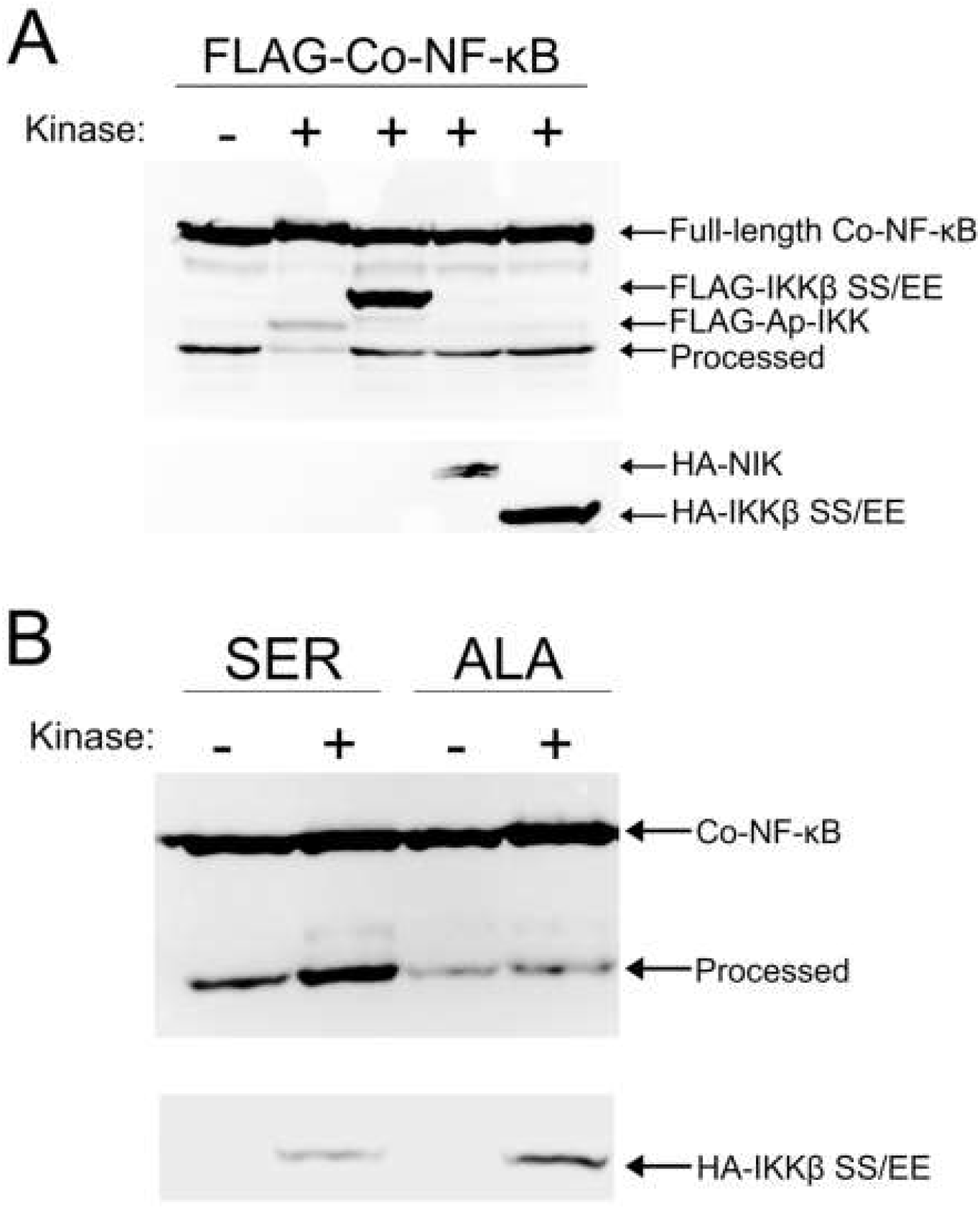
IKK-mediated processing of NF-κB arose with multicellularity. (A) Co-transfection with various kinases does not induce processing of FLAG-*Co*-NF-κB in HEK 293T cells. Arrows indicate the various FLAG- or HA-tagged kinases used in these assays. Full-length *Co*-NF-κB and processed *Co*-NF-κB are also indicated. (B) *Co*-NF-κB with C-terminal IKK target serines from *Aiptasia* NF-κB (*Co*-NF-κB-SER) or serine-to-alanine mutants (*Co*-NF-κB-ALA) were co-transfected with constitutively active human HA-IKKβ (SS/EE). Transfecting *Co*-NF-κB-SER and HA-IKKβ (SS/EE) resulted in the appearance of an increased amount of the lower band, but the alanine mutations (*Co*-NF-κB-ALA) abolished that processing.

To determine whether *Co*-NF-κB *could be* processed by an IKK-dependent mechanism, we created a mutant in which we replaced C-terminal sequences *Co*-NF-κB (downstream of the ANK repeats) with C-terminal sequences of the sea anemone *Aiptasia* (*Ap*)-NF-κB that contain conserved serines which can facilitate IKK-induced processing of *Ap*-NF-κB (17). We termed this mutant *Co*-NF-κB-SER, and also created the analogous protein (*Co*-NF-κB-ALA) in which the relevant serines were replaced by alanines. Co-expression of *Co*-NF-κB-SER with constitutively active human IKKβ (IKKβ SS/EE) resulted in increased amounts of the lower band, which was not seen with *Co*-NF-κB-ALA (Fig. 3B). Thus, the *Co*-NF-κB protein (consisting of the RHD, GRR, and ANK repeats) can undergo IKK-induced processing if supplied with a C terminus containing the IKK target serine residues. However, the native *Co*-NF-κB protein does not appear to be susceptible to IKK-induced processing, which is also consistent with the lack of any IKK sequences in the genome of *Capsaspora*.

### Exogenously expressed full-length and truncated versions of *Co*-NF-κB localize primarily to the nucleus in *Capsaspora* cells

We were next interested in examining the subcellular localization of NF-κB in *Capsaspora* cells. For these experiments, we transfected *Capsaspora* cells with our FLAG-tagged *Co*-NF-κB constructs (*Co*-NF-κB, mutant *Co*-RHD, and mutant *Co*-Cterm, see Fig. 2D) and then performed anti-FLAG indirect immunofluorescence. Consistent with results seen in DF-1 chicken fibroblast cells (Fig. 2D), FLAG-*Co*-RHD and FLAG-*Co*-Cterm localize to the nucleus and cytoplasm, respectively (Fig. 4, middle and bottom rows, Supplemental Video 1). Surprisingly, full-length FLAG-*Co*-NF-κB also appeared to be fully nuclear, as judged by its co-localization with the Hoechst-stained nuclei (Fig. 4, top row, and Supplemental Fig. 2 and Supplemental Video 2).

**Fig. 4.**
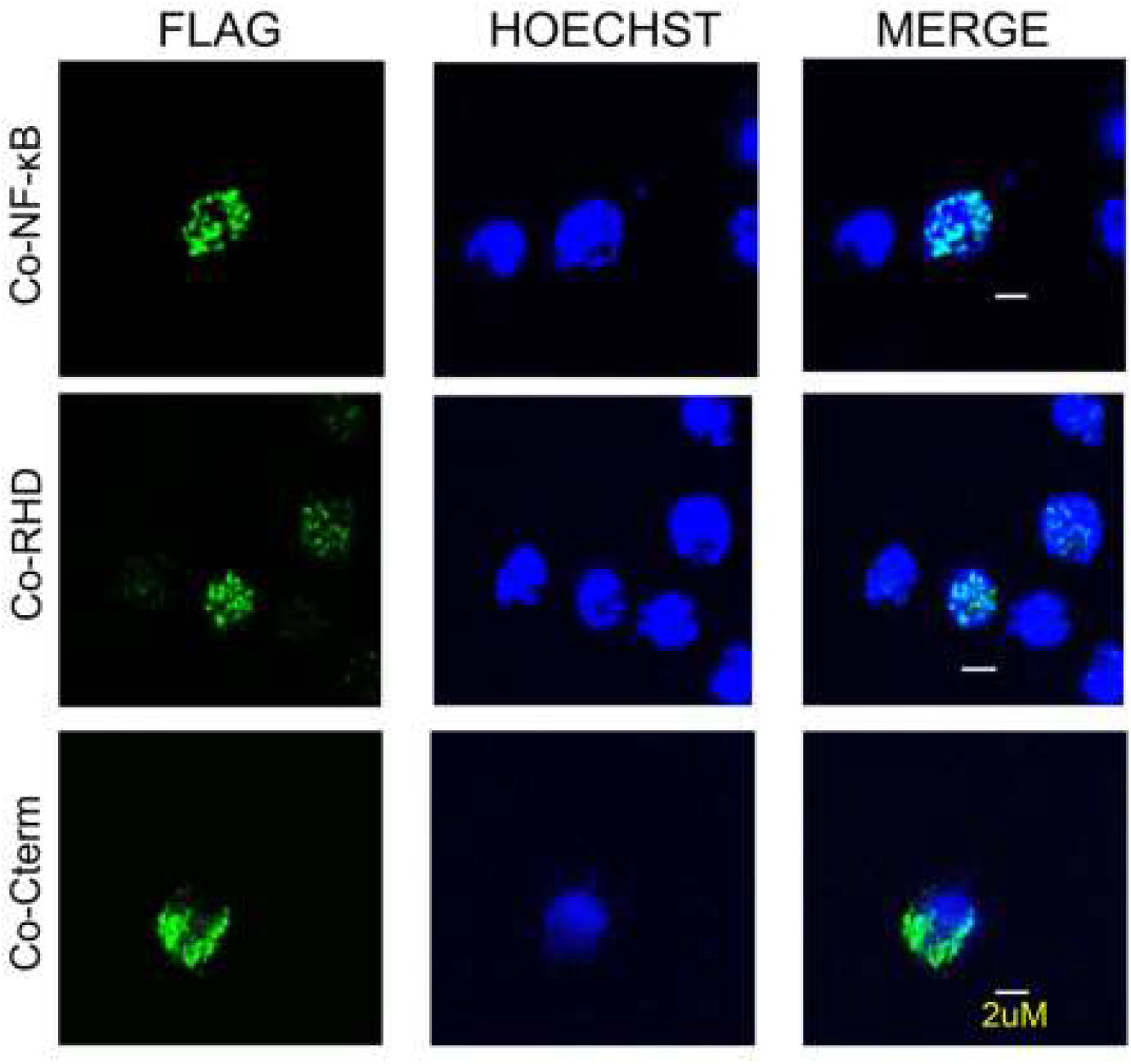
Transfection of FLAG-*Co*-NF-κB and FLAG-*Co*-RHD into *Capsaspora* cells results in nuclear localization. *Capsaspora* cells were transfected with FLAG-tagged vectors for full-length *Co*-NF-κB, mutant *Co*-RHD, and mutant *Co*-Cterm. The cells were stained using anti-FLAG antiserum (left panels) and HOECHST (middle panels), and then MERGED (right panels). Scale bars are 2 μM.

### *Co*-NF-κB mRNA levels and DNA-binding activity vary coordinately across different life stages and the identification of putative NF-κB target genes

*Capsaspora* has been shown to have three different life stages: aggregative, filopodia, and cystic, and RNA-Seq of each life stage has been reported (9). We were interested in whether *Capsaspora* NF-κB protein was active at different levels in these three life stages and whether we could use the previous RNA-Seq data to identify genes whose expression may be controlled by NF-κB. We first examined previous mRNA expression data (9) for NF-κB mRNA, and found that NF-κB was expressed at the lowest level in the aggregative stage and 2.3-and 5-fold higher in the filopodic and cystic stages, respectively (Fig. 5A). We next generated cultures of *Capsaspora* at each life stage (Fig. 5A), made protein extracts, and performed an EMSA using the κB-site probe that we showed can be bound by *Co*-NF-κB expressed in HEK 293T cells (Fig. 2E) and by bacterially expressed *Co*-RHD in our PBM assays (Supplemental Table 2). Consistent with the mRNA expression data, the κB-site probe was bound progressively stronger in aggregative, filopodic, and cystic stages (Fig. 5B). To determine whether the EMSA band indeed included *Co*-NF-κB, we incubated our protein extracts with 10X and 25X excesses of unlabeled κB-site probe, and we saw a substantial decrease in binding in the putative NF-κB band. In contrast, incubation with a 25X excess of an unlabeled IRF-site probe did not decrease the putative *Co*-NF-κB complex, indicating that the binding was specific for the κB-site probe.

**Fig. 5.**
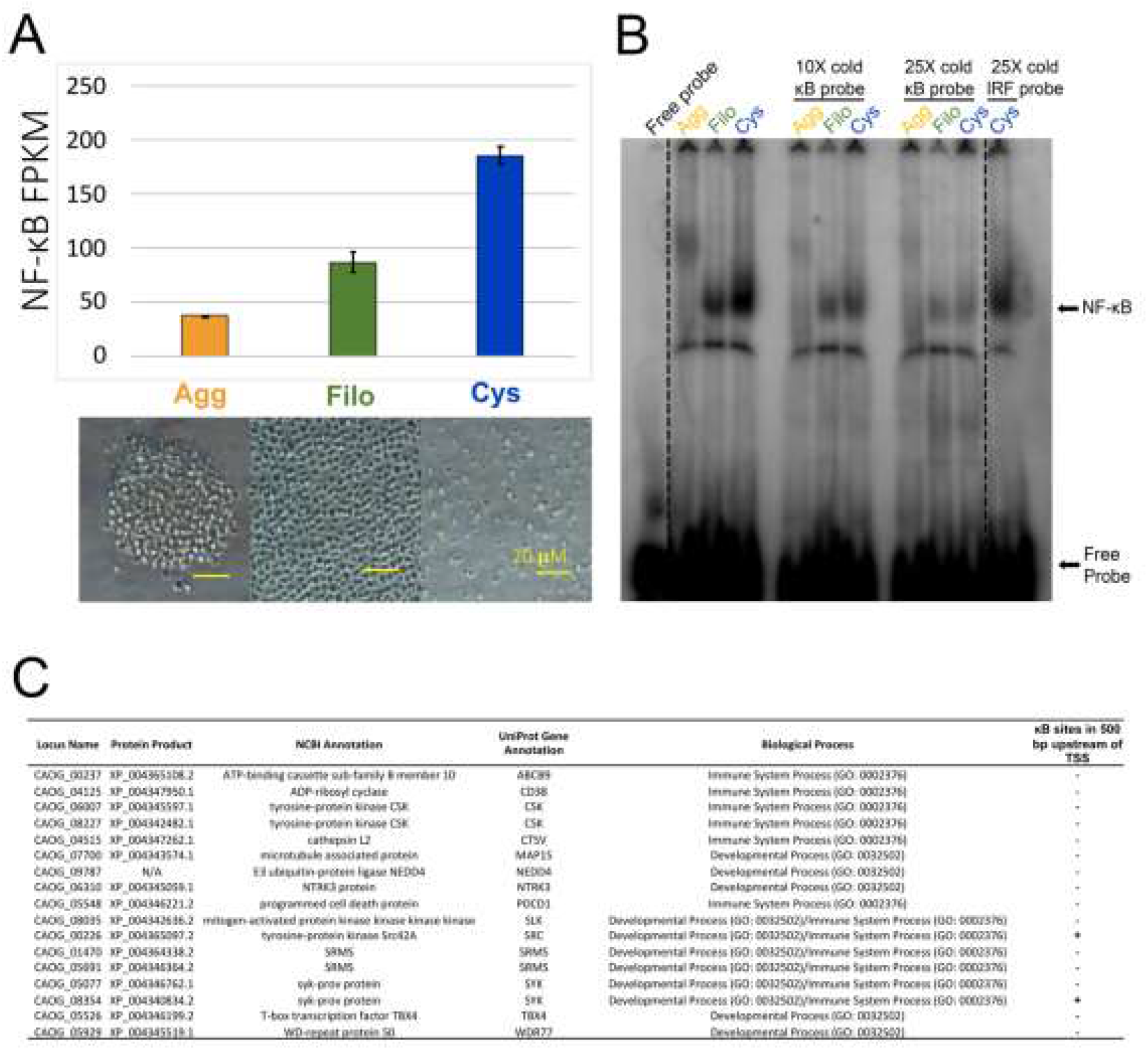
NF-κB is differentially expressed during the different life stages of *Capsaspora* and its proposed roles in development and immunity. (A) Top: The FPKM values from Sebé-Pedrós et al. (9) of NF-κB at each life stage, done in triplicate. Agg, Aggregative (yellow), Filo, Filopodic (green), Cys, Cystic (blue). Error bars are standard deviation. Bottom: Images taken with a light microscope of each life stage (Agg, Filo, and Cys from left to right). Yellow scale bar is 20 μM. (B) *Capsaspora* whole-cell extracts were created from each life stage (see Methods). 70 μg of each extract was then used in an electromobility shift assay (EMSA). Lane 1 contains only free probe.Lanes 2-4 contain lysates from Agg, Filo, and Cys life stages incubated with a radioactive κB-site probe. Lanes 5 and 9 are empty. Lanes 6-8, and lanes 10-12 contain lysates from Agg, Filo, and Cys life stages as indicated, and were incubated with an excess (10X and 25X, respectively) of unlabeled κB-site probe. Lane 13 contains the Cys lysate incubated with 25X unlabeled IRF-site probe. NF-κB complexes and Free Probe are indicated with arrows. The dashed lines indicate where the gel was cut to remove excess lanes. (C) NF-κB expression may influence the expression of genes that are involved in developmental and immune system processes. The expression profiles of these genes correlate with NF-κB mRNA expression in each life stage (Agg, low; Filo, medium; Cys, high), and were identified via Biological Processes GO analysis. Two of the genes in this list (SYK and SRC) also contain κB sites in the 500 bp upstream of their TSS.

We next sought to identify genes that might be influenced by the expression of NF-κB in order to gain insight into potential functional roles for NF-κB in these life stages. We examined the existing RNA-Seq data (9), which contains the mRNA expression of 8674 genes of *Capsaspora* during its three life stages. We first narrowed our gene list to those genes that were differentially expressed in a manner similar to NF-κB mRNA levels and DNA-binding activity during each life stage (i.e., progressively increased in expression from aggregative, filopodic, and cystic stages). From this exercise, we identified 1348 mRNAs that were expressed at low levels in the aggregative stage, and successively higher levels in the filopodic and cystic stages.

Of the 1348 genes that we identified, 389 genes were annotated (which is consistent with approximately 1/4 of *Capsaspora’s* total predicted protein-encoding genes being annotated, Supplemental Table 3), and 305 of these genes had human homologs (Supplemental Table 3). We then performed GO analysis to look at Biological Processes overrepresented in these 305 genes. That analysis showed that this set of 305 genes was predicted to be involved in several biological processes, including 14 genes that encode proteins associated with developmental and immune system processes (Fig. 5C), which are biological processes regulated by NF-κB in many more complex organisms and suggested to be regulated by NF-κB in several basal organisms (15–17, 20–23). Although 14 genes may seem low, the total database for human GO analysis of immune system and developmental processes genes is approximately 2200 genes, but the number of annotated homologs that exist in *Capsaspora* in these two GO categories is only 66 genes (Supplemental Table 4). Thus, about 20% (14/66) of the *Capsaspora* genes in the GO category for the developmental and immune processes subset are among the 305 homologous genes coordinately regulated with NF-κB. Other broad categories overrepresented in these 305 genes included Signaling, Metabolic Process, and Locomotion (Supplemental Fig. 3).

We then looked for κB sites within 500 bp upstream of the transcription start sites (TSS) for each of the 1348 genes with expression profiles that were similar to NF-κB. 192 of these 1348 gene upstream regions contained 1-3 κB sites within 500 bp of the TSS, with the majority of these genes containing 1 κB site (Supplemental Table 5). Two of the 14 genes that encode protein homologs associated with GO developmental and immune system processes contained a κB site within the 500 bp upstream of their TSS (Fig. 5C).

### Choanoflagellate NF-κBs can form heterodimers and have different abilities to bind DNA and activate transcription

Richter et al. (5) sequenced the transcriptomes of 19 choanoflagellates and identified RHD-containing NF-κB-related proteins in 12 of these species. We chose to characterize the NF-κB proteins from *Acanthoeca spectabilis* (*As*) because it has three NF-κB-like proteins, which separated into multiple branches when phylogenetically compared to all choanoflagellate NF-κBs (Fig. 1B). These three *As*-NF-κB proteins contained ostensibly complete DNA-binding sequences, which were similar to other NF-κB proteins and a putative NLS (3). These three proteins contained extended sequences N-terminal to the RHD, but they contained few C-terminal residues beyond the RHD (and no GRRs or ANK repeats). As a first step in characterizing these proteins, we created pcDNA FLAG vectors for *As*-NF-κB1, *As*-NF-κB2, and *As*-NF-κB3 (Fig. 6A) and transfected them into HEK 293T cells. As assessed by anti-FLAG Western blotting, each vector directed the expression of appropriately sized FLAG-tagged proteins (Fig. 6B).

**Fig. 6.**
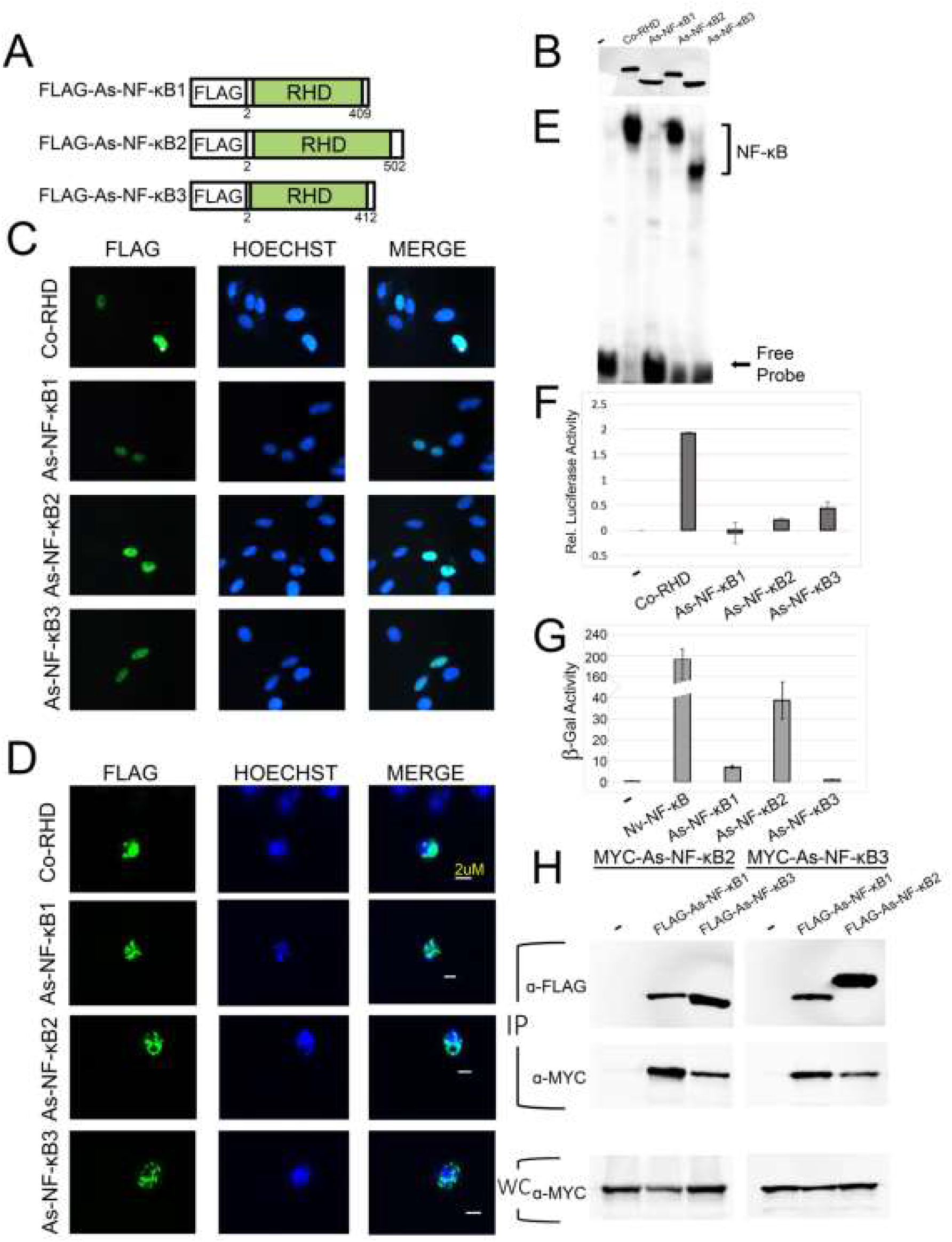
Characterization of cellular and molecular properties of three choanoflagellate NF-κBs. (A) FLAG-tagged NF-κB proteins used in these experiments. From top to bottom the drawings depict the three NF-κB-like proteins from the transcriptome of *A. spectabilis* (5). RHDs are in green (Rel Homology Domain). (B) Anti-FLAG Western blot of lysates from HEK 293T cells transfected with the indicated expression vectors. (C and D) Indirect immunofluorescence of DF-1 chicken cells (C) or *Capsaspora* cells (D) transfected with the indicated expression vectors. Cells were then stained with anti-FLAG antiserum (left panels) and HOESCHT (middle panels), and then MERGED in the right panels. Yellow scale bar in *Capsaspora* panels is 2 μM. (E) A κB-site electromobility shift assay (EMSA) using each of the indicated lysates from (B). The NF-κB complexes and free probe are indicated by arrows. (F) A κB-site luciferase reporter gene assay was performed with the indicated proteins in HEK 293 cells. Luciferase activity is relative (Rel.) to that seen with the empty vector control (1.0), and values are averages of three assays performed with triplicate samples with standard error. Values are shown on a log scale. (G) A GAL4-site *LacZ* reporter gene assay was performed with GAL4-fusion proteins in yeast Y190 cells. Values are average units of two assays performed with four samples with standard error. (H) Coimmunoprecipitation (IP) assays of MYC-tagged *As*-NF-κB2 and *As*-NF-κB3. In each IP assay, MYC-*As*-NF-κBs were co-transfected with pcDNA FLAG, FLAG-*As*-NF-κB1, 2 or 3 as indicated. An IP using anti-FLAG beads was performed. Anti-FLAG (top) and anti-MYC (middle) Western blotting was then performed. An anti-MYC Western blot of the whole-cell (WC) lysates was also performed (bottom).

To determine the subcellular localization properties of the three *As*-NF-κBs, we performed indirect immunofluorescence on DF-1 chicken fibroblasts and *Capsaspora* cells transfected with each FLAG-tagged vector. All three *As*-NF-κB proteins co-localized with Hoechst-stained nuclei in both DF-1 cells (Fig. 6C) and *Capsaspora* cells (Fig. 6D).

We then performed a κB-site EMSA on whole cell extracts from HEK 293T cells overexpressing each *As*-NF-κB, using *Co*-RHD as a positive control. *As*-NF-κB2 and 3 bound the κB-site probe to nearly the same extent as *Co*-RHD, but *As*-NF-κB1 only weakly bound the probe (Fig. 6E). We also assessed the transactivating ability of each *As*-NF-κB protein in a κB-site reporter assay in HEK 293T cells, using the strongly activating *Co*-RHD protein as a positive control. Both *As*-NF-κB2 and 3 were able to activate transcription of the luciferase reporter above vector control levels (~1.6- and 2.7-fold, respectively) but *As*-NF-κB1 did not (Fig. 6F). We also assessed the intrinsic transactivating ability of each *As*-NF-κB protein using a GAL4-fusion reporter assay in yeast cells. In this assay, all three *As*-NF-κBs activated transcription over vector control levels, although *As*-NF-κB1 and 3 activated to a much lesser degree than *As*-NF-κB2 (Fig. 6G). From these data, we hypothesized that the homodimeric version of *As*-NF-κB1 was not capable of binding DNA and could not activate transcription using κB sites, but likely contained some intrinsic ability to activate transcription (as a GAL4-fusion protein), whereas homodimeric *As*-NF-κB2 and 3 could activate transcription both in human cell-based and yeast GAL4-fusion reporter assays.

Since *As*-NF-κB1 did not substantially bind DNA or activate transcription when transfected alone, we hypothesized that *As*-NF-κB1 ordinarily acts as a heterodimer with the other *As*-NF-κBs. Therefore, we performed a series of co-immunoprecipitation experiments to determine whether *As*-NF-κB1 could interact with *As*-NF-κB2 or *As*-NF-κB3. To do this, we subcloned *As*-NF-κB2 and 3 into MYC-tagged vectors, co-transfected each with FLAG-*As*-NF-κB1 in HEK 293T cells, and first immunoprecipitated cell extracts using anti-FLAG beads. We then performed anti-FLAG and anti-MYC Western blotting on the immunoprecipitates to assess whether these NF-κBs could interact. MYC-*As*-NF-κB2 and MYC-*As*-NF-κB3 were both co-immunoprecipitated with FLAG-*As*-NF-κB1, as well as with each other (Fig. 6H, IP). The MYC-*As*-NF-κB proteins were not seen when they were co-transfected with the empty vector control (Fig. 6H, IP).

From these data, it appears that all three *As*-NF-κBs can enter the nucleus when expressed in vertebrate and protist cells, but they bind DNA and activate transcription to varying degrees. Furthermore, all three *As*-NF-κBs can form heterodimers with the other *As*-NF-κBs. As we discuss below, we think that the reduced ability of *As*-NF-κB1 to bind DNA and activate a reporter gene is due to a limited ability to form homodimers.

## Discussion

In this manuscript, we have functionally characterized and compared, for the first time, NF-κB proteins from two protists. Taken together, these results demonstrate that although functional DNA-binding and transcriptional-activating NF-κB proteins exist in these protists, the overall structures and regulation of these proteins varies considerably, both among protists and when compared to animal NF-κBs.

In the vertebrate NF-κB proteins p100 and p105, C-terminal ANK repeats inhibit the DNA-binding activity of the RHD, and proteasome-mediated processing of the ANK repeats is terminated by and within the GRR (13). In the *Drosophila* Relish protein, the C-terminal ANK repeats also inhibit DNA binding, but the C-terminal ANK repeats are removed by an internal site-specific cleavage event, which does not involve the proteasome, and Relish has no GRR (24). Thus, the presence of ANK repeats and a GRR in *Co*-NF-κB suggests that proteasomal processing would lead to nuclear translocation and activation of its DNA-binding activity. Indeed, removal of the C-terminal residues of *Co*-NF-κB does allow it to enter the nucleus of vertebrate cells and unleashes its DNA-binding activity (Figs. 2D and E). Moreover, treatment of cells with the proteasome inhibitor MG132 blocks the basal processing of *Co*-NF-κB that is seen in transfected 293T cells (Supplemental Fig. 1). However, this basal processing of *Co*-NF-κB in 293T cells is not enhanced by co-expression of IKK (unless C-terminal IKK target sequences are added, see Fig. 3B) and there are no obvious IKK target serines in *Co*-NF-κB nor are there any IKK homologues in the *Capsaspora* genome. Furthermore, most of the *Co*-NF-κB appears to be in the nucleus when it is overexpressed in *Capsaspora* cells (Fig. 4), suggesting that it is constitutively processed to its RHD. If *Co*-NF-κB does undergo a signal-induced proteasomal processing in *Capsaspora* cells, then it is unlikely to be dependent on an IKK-like kinase. Alternatively, but probably less likely, full-length *Co*-NF-κB enters the nucleus of *Capsaspora* cells but not DF-1 chicken cells. In the absence of a *Co*-NF-κB-specific antiserum, we cannot distinguish between these possibilities.

In contrast to *Capsaspora*, the choanoflagellate NF-κB proteins lack C-terminal ANK repeats and GRRs, and all three *As*-NF-κB proteins are constitutively in the nucleus when overexpressed in vertebrate or *Capsaspora* cells (Fig. 6). We have not been able to identify an IκB-like protein in the *A. spectabilis* genome. Thus, it is unclear whether choanoflagellate NF-κB is regulated by an ANK-dependent cytoplasmic retention mechanism, or whether, for example, choanoflagellate proteins are constitutively nuclear in their native settings. Nevertheless, it is clear that the regulation of both *Capsaspora* and choanoflagellate NF-κB proteins is distinct from what is seen with NF-κB proteins in higher metazoans.

Of interest, constitutively nuclear localization of NF-κB proteins has also been seen in other settings. That is, we have previously shown that in both the sea anemone *Aiptasia* and sponges, most NF-κB staining is nuclear and the proteins are mostly processed in their native settings (15–17). Thus, we have argued previously (3) that these basal NF-κB proteins may be constitutively in an active state, perhaps due to continual interaction with upstream activating ligands or pathogens. Of note, most NF-κB p100 is also in its processed form in mouse spleen tissue (25).

Among the 21 choanoflagellates for which there is sufficient transcriptomic/genomic information, it appears that only 12 have any NF-κB genes. Surprisingly, in seven of these 12 choanoflagellates there are multiple NF-κB genes. Thus, it is clear that there have been gains and losses of NF-κB genes among the choanoflagellates. We note that NF-κB has also been lost in other organisms including *C. elegans* and ctenophores (14). The absence of NF-κB in some choanoflagellates and its expansion in others (e.g., *A. spectabilis*) suggests that NF-κB has a specialized, rather than a general, biological function in choanoflagellates.

The presence of three NF-κB-like heterodimerizing proteins in *A. spectabilis* is the first example of an organism basal to flies with multiple NF-κB family proteins that are capable of forming heterodimers. Thus, expansion of NF-κB genes has occurred multiple times in evolution, i.e., at least once in the metazoan lineage and once within choanoflagellates. Furthermore, since each *As*-NF-κB homodimer has a different ability to bind DNA and activate transcription, it appears that there are subclasses of NF-κB within *A. spectabilis* and likely within other choanoflagellates that have multiple NF-κBs. It is interesting to note that *As*-NF-κB1, 2, and 3 are phylogenetically separate and cluster most closely with NF-κBs from certain other choanoflagellate species that contain multiple NF-κBs. For example, *Savillea parva* contains three NF-κBs, each of which clusters with a separate *As*-NF-κB (Fig. 1B). Thus, we hypothesize that choanoflagellates, like vertebrates, have evolved a mechanism for differential transcriptional control of genes through the use of combinatorial dimer formation.

The differential mRNA expression and DNA-binding activity of NF-κB among different life stages of *Capsaspora* suggest that NF-κB has life stage-specific functions. It is notable that the DNA-binding activity of NF-κB in these different life stages correlates with differences in the levels of NF-κB mRNA, rather than as differences in induced activity. That is, in most metazoans, the activity of NF-κB is regulated at the post-transcriptional level, whereas in *Aiptasia* and corals, we have found that NF-κB mRNA levels and DNA-binding activity appear to be coordinately regulated, suggesting transcriptional regulation. That is, in *Aiptasia*, thermal bleaching causes transcriptional upregulation of NF-κB, which also results in increased protein expression of nuclear, DNA binding-active NF-κB (17), which is similar to what we see with NF-κB across the *Capsaspora* life stages. Similarly, treatment of the coral *Orbicella faveolata* with lipopolysaccharide results in increased expression of NF-κB target genes, rather than increased post-translational activation of NF-κB (15). Thus, it appears that in several basal organisms NF-κB proteins are constitutively nuclear and that increases in their activity is the result of transcriptional upregulation of NF-κB mRNA, rather than induced proteolysis, which occurs in most mammalian and fly systems.

Activation of NF-κB by signal-induced degradation of IκB sequences is essentially dogma in vertebrates. Although several basal NF-κBs, including *Co*-NF-κB, can be formulated to undergo IKK-induced processing when expressed in human cells in culture (15–17), there is much evidence that such regulation may not occur in the native animals. For example, although sponge and sea anemone *Aiptasia* NF-κBs have C-terminal ANK repeats and GRRs, they are largely processed and nuclear in the animals themselves (15–17). Moreover, induction of NF-κB DNA-binding activity and protein levels by loss of symbiosis in *Aiptasia* appears to be primarily a result of increased transcription of NF-κB and not due to induced ANK repeat degradation. Similarly, the increased NF-κB DNA-binding activity seen in different life stages of *Capsaspora* is paralleled by increased expression of NF-κB mRNA in these life stages. Furthermore, when *Co*-NF-κB is overexpressed in *Capsaspora* cells it is primarily in the nucleus, likely due to constitutive processing to the RHD form. Thus, a more relevant question for many basal NF-κB proteins may be what conditions stop them from being processed. Of note, the three *As*-NF-κBs do not have C-terminal ANK repeats and we have not been able to identify a putative IκB in *A. spectabilis* transcriptomic databases. Therefore, it is possible that the activity of these constitutively nuclear choanoflagellate NF-κBs is fully regulated by transcriptional control of their genes.

Of the nearly 1350 genes whose expression correlated with NF-κB expression across different *Capsaspora* life stages, almost 20% contained κB binding sites within 500 base pairs upstream their TSS (Fig. 5). While this might be an overestimate of NF-κB gene targets or genes indirectly influenced by expression of NF-κB, there are likely additional NF-κB binding sites that could affect target gene expression. For example, ATAC-seq data have suggested that the regulatory sites in the *Capsaspora* genome are present in first introns, 5’ UTRs, as well as the proximal intergenic regions (26). Among the ~1350 genes that we identified with expression patterns similar to NF-κB, there are 192 genes that contain NF-κB binding sites in their proximal promoters. Two of these genes are homologs of SYK and SRC (CAOG_08354 and CAOG_00206), which are known in higher organisms to be involved in immunity and development. However, the list of ~1350 genes most certainly contains genes that are controlled by other transcription factors or are regulated by signaling or developmental events that are partially or not at all affected by NF-κB. Nevertheless, it is clear that NF-κB might play several roles, and perhaps different roles, in each life stage of *Capsaspora*.

Our studies suggest that the regulation and associated biology of NF-κB in single-celled organisms are different from what is seen in multicellular vertebrates and flies. Perhaps, the concerted effort of aggregation in *Capsaspora* and the correlative decrease in NF-κB in these aggregated cells reflect a need to suppress collective immunity to form a symbiotic group. Alternatively, *Capsaspora* is normally a symbiont in the hemolymph of the snail *B. glabrata*, where NF-κB may play a role in maintaining symbiosis, which has been suggested as one function of NF-κB in other organisms (27).

It is not clear what type of pathway might lead to activation of NF-κB in protists. In animals across a broad swath of phyla, the binding of a ligand to receptors such as TLRs/IL-1Rs or TNFRs initiates signaling pathways that converge on an IKK complex which then activates NF-κB (1). However, in basal organisms, many of these components are missing, few in number, or lack critical domains (Supplemental Fig. 4). For example, while some cnidarians contain homologs to TLRs, other cnidarians, some sponges, and choanoflagellates contain only TIR-domain proteins that lack the important extracellular components of the TLR (5, 11, 15, 16, 28). Furthermore, *Capsaspora* does not contain any homologs to TLRs, ILR-1 or TNFRs (4).

The diversification in NF-κB that we see here between *Capsaspora* and choanoflagellates, members of the same supergroup of protists, suggests that the diversification of NF-κB among all protists will be considerable. Overall, our results contribute to an understanding of NF-κB across extant phyla. The continued study of the evolution of NF-κB and other basally derived transcription factors will likely lead to an understanding of where and how these factors originated, as well as the basal biological functions they control.

## Materials and methods

### Phylogenetic analyses

The RHD sequences of NF-κB from *C. owczarzaki* were compared phylogenetically to the NF-κB-like sequences present in the transcriptomes of sequenced choanoflagellates. Details on databases and sequence acquisition can be found in Supplemental Table 6. Sequences were aligned by Clustal Omega (29). A maximum likelihood phylogenetic tree was created using PAUP* (30) and was bootstrapped 1000 times. Bayesian phylogenetic analyses showed similar results (data not shown).

### Plasmids

Expression plasmids for FLAG-tagged human IKKβ, FLAG-Nv-NF-κB, HA-Hu-IKKβ– SS/EE (constitutively active), FLAG-Ap-IKK, and the empty pcDNA-FLAG vector have been described previously (17, 31). The cDNA of the codon-optimized NF-κB sequence of *Capsaspora* was synthesized in a pUC57 plasmid by GenScript and was then subcloned into pcDNA-FLAG. PCR-generated Co-NF-κB truncation mutants (Co-RHD and Co-Cterm) were subcloned into pcDNA-FLAG or the yeast GAL4-fusion vector pGBT9. Three human codon-optimized *As*-NF-κB (named 1, 2, and 3) cDNAs were synthesized by GenScript based on sequences from the transcriptome of *A. spectabilis*. These cDNAs were subcloned into pcDNA-FLAG. To create MYC-tagged versions of As-NF-κB2 and 3, the sequences were excised from their pUC57 vectors using EcoRI and XhoI, and were then subcloned into a MYC-tagged expression vector (a gift of Shigeki Miyamoto, University of Wisconsin). Details of primers and plasmid constructions can be found in Supplemental Tables 7 and 8, respectively.

### *Capsaspora* culture, transfection, and protein extraction

*Capsaspora* cell cultures (strain ATCC ®30864) were grown axenically in 25 cm^2^ culture flasks (ThermoScientific Nunclon) with 10 ml ATCC medium 1034 (modified PYNFH medium) at 23°C. Cultures for each *Capsaspora* life stage were generated as previously described (9) and as instructed by ATCC. Filopodic cells were maintained in an actively dividing adherent state by scraping and passaging 1/40-1/50 of the cultures every 6-8 days, before floating cells appeared. Floating cystic cells were collected from 14 day-old filopodic cultures. Aggregative cells were created by actively scraping dividing filopodic cells and seeding them into a 25 cm^2^ culture flask, which was gently agitated at 60 RPM for 4-5 days, and grown axenically at 23°C. Whole-cell lysates of cells from each life stage were prepared in AT Lysis Buffer (20 mM HEPES, pH 7.9, 150 mM NaCl, 1 mM EDTA, 1 mM EGTA, 20% w/v glycerol, 1% w/v Triton X-100, 20 mM NaF, 1 mM Na_4_P_2_O_7_·10H_2_O, 1 mM dithiothreitol, 1 mM phenylmethylsulfonyl fluoride, 1 μg/ml leupeptin, 1 μg/ml pepstatin A, 10 μg/ml aprotinin), as described previously (19).

Transfection of *Capsaspora* cells with expression plasmids was performed using polyethylenimine (PEI) (Polysciences, Inc.). Briefly, the day before transfection, actively dividing *Capsaspora* filopodic cells were plated at about 80-90% confluency. The next day, cells were transfected by incubation with 5 μg of plasmid DNA and 25 μl of 1 mg/ml PEI. Media was changed ~20 h post-transfection, and *Capsaspora* cells to be analyzed by immunofluorescence were passaged onto poly-D-lysine (ThermoFisher)-treated glass coverslips on the day prior to fixation.

### Vertebrate cell culture and transfection

DF-1 chicken fibroblasts and human HEK 293 or 293T cells were grown in Dulbecco’s modified Eagle’s Medium (DMEM) (Invitrogen) supplemented with 10% fetal bovine serum (Biologos), 50 units/ml penicillin, and 50 μg/ml streptomycin as described previously (19). Transfection of cells with expression plasmids was performed using polyethylenimine (PEI) (Polysciences, Inc.) as described previously (19, 31). Briefly, on the day of transfection, cells were incubated with plasmid DNA and PEI at a DNA:PEI ratio of 1:6. Media was changed ~20 h post-transfection, and whole-cell lysates were prepared 24 h later in AT Lysis Buffer (20 mM HEPES, pH 7.9, 150 mM NaCl, 1 mM EDTA, 1 mM EGTA, 20% w/v glycerol, 1% w/v Triton X-100, 20 mM NaF, 1 mM Na_4_P_2_O_7_·10H_2_O, 1 mM dithiothreitol, 1 mM phenylmethylsulfonyl fluoride, 1 μg/ml leupeptin, 1 μg/ml pepstatin A, 10 μg/ml aprotinin). DF-1 cells analyzed by immunofluorescence were passaged onto glass coverslips on the day prior to fixation.

### Western blotting, electrophoretic mobility shift assays (EMSAs), reporter gene assays, and indirect immunofluorescence

Western blotting was performed as described previously (17, 19). Briefly, cell extracts were first separated on 7.5% or 10% SDS-polyacrylamide gels. Proteins were then transferred to nitrocellulose at 4°C at 250 mA for 2 h followed by 170 mA overnight. The membrane was blocked in TBST (10 mM Tris-HCl [pH 7.4], 150 mM NaCl, 0.1% v/v Tween 20) containing 5% powered milk (Carnation) for 1 h at room temperature. Filters were incubated at 4 °C with FLAG primary antiserum (1:1000, Cell Signaling Technology) diluted in 5% milk TBST. After extensive washing in TBST, filters were incubated with anti-rabbit horseradish peroxidase-linked secondary antiserum (1: 4000, Cell Signaling Technology). Immunoreactive proteins were detected with SuperSignal West Dura Extended Duration Substrate (Pierce) and imaged on a Sapphire Biomolecular Imager (Azure Biosystems).

EMSAs were performed using a ^32^P-labeled κB-site probe (GGGAATTCCC, see Supplemental Table 6) and 293T or *Capsaspora* whole-cell extracts prepared in AT buffer (17, 19). EMSA gels were exposed on a phosphor screen, and then imaged on a Sapphire Biomolecular Imager (Azure Biosystems). Yeast GAL4-site *LacZ* and 293 cell κB-site luciferase reporter gene assays were performed as described previously (19). Transfection and indirect immunofluorescence of DF-1 and *Capsaspora* cells were performed on methanol-fixed or paraformaldehyde-fixed, respectively, cells that were probed with rabbit anti-FLAG primary antiserum (1:80, Cell Signaling Technology), essentially as described previously (19). Nuclei were stained with Hoechst as indicated.

### RNA-sequencing analysis and annotation

RNA-sequencing data from each life stage of *Capsaspora* was obtained from Sebé-Pedrós et al. (9). To identify potential target genes of NF-κB, we sorted all sequencing data with genes that were differentially expressed in the same manner as NF-κB for each life stage (lowest expression in aggregative, medium expression in filopodic, highest expression in cystic). We discarded genes if they were not expressed at a life stage (i.e., had an RPKM of 0). Of the 8674 total genes, 1348 genes were differentially expressed in a manner similar to NF-κB. Of these 1348 genes, 389 genes had annotated homologs, and 305 had human homologs (Supplemental Table 3). We then performed GO analysis (http://pantherdb.org) by entering the UniProt ID of each gene and selecting *Homo sapiens*. We then created and analyzed the Biological Processes that were present in this gene list.

To identify genes with potential upstream NF-κB binding sites among these 1348 genes, we also extracted the 500 base pair sequence upstream of each gene. We then imported these 1348 upstream regions into MEME-FIMO and scanned the sequences for NF-κB motifs (but not Rel motifs) extracted from JASPAR (http://jaspar.genereg.net) (Supplemental Table 4).

### Immunoprecipitations

HEK 293T cells were transfected with MYC-As-NF-κB2 and either a pcDNA FLAG empty vector, FLAG-As-NF-κB1, or FLAG-As-NF-κB3 as above (See “Vertebrate cell culture and transfection”). Lysates were prepared 48 h later and were incubated with 50 μl of a 1X PBS-washed anti-FLAG bead slurry (Sigma) overnight at 4 °C with gentle rocking. The next day, the beads were washed three times with 1X PBS. The pellet was then boiled in 2X SDS sample buffer, and the supernatant was electrophoresed on a 7.5% SDS-polyacrylamide gel, as above (see “Western blotting”). The membrane was then probed with a rabbit anti-MYC (Cell Signaling Technologies, 1:1000) antiserum, then with anti-rabbit horseradish peroxidase-linked secondary antiserum (1: 4000, Cell Signaling Technology), and reacted with SuperSignal West Dura substrate (Pierce). An image was then obtained on a Sapphire Biomolecular Imager (Azure Biosystems). The membrane was stripped and probed with rabbit anti-FLAG antiserum as above. These same transfection and IP experiments were repeated with MYC-As-NF-κB3 and either pcDNA FLAG empty vector, FLAG-As-NF-κB1, or FLAG-As-NF-κB2.

## Acknowledgments

This research was supported by following National Science Foundation grants (to T.D.G.): IOS-1354935 and IOS-1937650. L.M.W. was supported by an NSF Graduate Research Fellowship and a Warren-McLeod Graduate Fellowship in Marine Sciences (Boston University). J.S. and E.K.A. were supported by NSF REU BIO-1659605 (T.D.G.). S.S. was supported by the Boston University (BU) Undergraduate Research Opportunities Program, and A.A. was supported by the BU GROW program. Designated authors performed research as part of the undergraduate Molecular Biology Laboratory course BB522 (Spring, 2019, 2020), and were supported by funds from the BU Biology Department.

## Competing Interest Statement

We have no competing interests.

## Author Contributions

L.M.W. and T.D.G planned the project and wrote the manuscript, and L.M.W performed experiments except those that follow. S.S. cloned plasmids used in Fig. 6 and performed Western blot in Sup. Fig. 1. J.S. and E.A. did initial Western blots for Fig. 3 and Luciferase assay on Fig. 2. A.A. analyzed and selected As-NF-κBs. BB522 Molecular Biology Laboratory, N.R-S. and C.J.D. created plasmids for Figs. 2B and 6A, and C.J.D. took images for Fig. 6B. P.J.A.C. performed EMSA on Fig. 2E. T.S. analyzed PBMs in Fig. 2A.

## Notes

### Competing Interest Statement

The authors have declared no competing interest.

### Summary of Updates

We have changed the NSF grant support in Acknowledgments, since it came to our attention that it was cited incorrectly in the previous version. Nothing else in the manuscript has changed.

